# Oxygen affinities of DosT and DosS sensor kinases with implications for hypoxia adaptation in *Mycobacterium tuberculosis*

**DOI:** 10.1101/2024.02.26.582189

**Authors:** Elizabeth A. Apiche, Eaindra Yee, Anoop Rama Damodaran, Ambika Bhagi-Damodaran

## Abstract

DosT and DosS are heme-based kinases involved in sensing and signaling O_2_ tension in the microenvironment of *Mycobacterium tuberculosis* (*Mtb*). Under conditions of low O_2_, they activate >50 dormancy-related genes and play a pivotal role in the induction of dormancy and associated drug resistance during tuberculosis infection. In this work, we reexamine the O_2_ binding affinities of DosT and DosS to show that their equilibrium dissociation constants are 3.3±1 μM and 0.46±0.08 μM respectively, which are six to eight-fold stronger than what has been widely referred to in literature. Furthermore, stopped-flow kinetic studies reveal association and dissociation rate constants of 0.84 μM^-1^s^-1^ and 2.8 s^-1^, respectively for DosT, and 7.2 μM^-1^s^-1^ and 3.3 s^-1^, respectively for DosS. Remarkably, these tighter O_2_ binding constants correlate with distinct stages of hypoxia-induced non-replicating persistence in the Wayne model of *Mtb*. This knowledge opens doors to deconvoluting the intricate interplay between hypoxia adaptation stages and the signal transduction capabilities of these important heme-based O_2_ sensors.

## Main Text

Heme-based sensors play a crucial role in enabling a wide range of microbes to detect and respond to changes in their redox environment [1,2]. Among these, DosT and DosS are two extensively investigated heme-based sensor kinases that play a vital role in the recognition and response mechanism of *Mycobacterium tuberculosis* (*Mtb*), the causative agent of tuberculosis (TB), to varying levels of O_2_ [3–6]. In low O_2_ environments, such as those found in TB granulomas, these sensors switch on their histidine kinase activity and initiate the expression of over 50 genes responsible for inducing a state of dormancy (**Fig. 1a**) [7]. The kinase activity of DosT and DosS is inhibited in their O_2_-bound form [8,9]. However, exposure to environments with low O_2_ causes these heme-based sensor kinases to lose the O_2_ molecule bound to their heme domain and switch to their active ferrous form, resulting in the autophosphorylation of specific amino acid residues, His 392 and His 395 in DosT and DosS, respectively [10]. Remarkably, these heme-based sensors remain active even upon binding to redox-active ligands such as NO and CO, that are present in TB granulomas [8,11]. The phosphorylated DosT and DosS subsequently transfer the phosphoryl group to the DosR response regulator, which then binds to DNA to activate dormancy transformation in *Mtb* [4]. Under dormancy, *Mtb*’s metabolism significantly slows down, rendering it non-replicative and simultaneously resistant to current antibiotic treatments [12].

**Figure 1.**
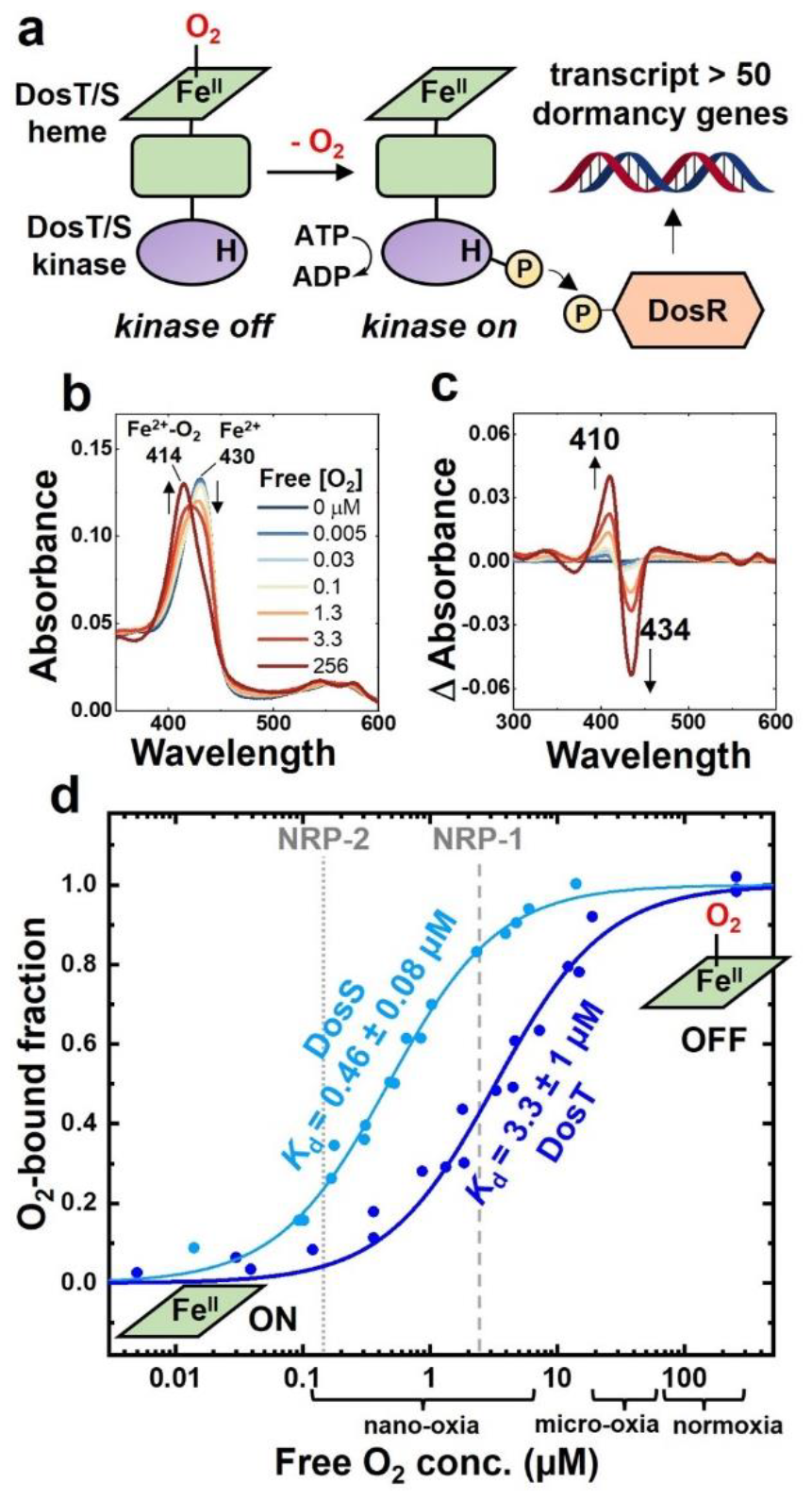
**a)** Function of DosT and DosS sensors in *Mtb*. In their O_2_ bound forms, the kinase domains of sensors are switched off. Under O_2_ tension, heme loses O_2_ to turn on the kinase domain and phosphorylate His 392 and 395 residues in DosT and DosS, respectively. Phosphorylated forms of DosT/S transfer the phosphoryl group to DosR which binds to *dosR* regulon and activates dormancy signaling in the bacteria. **b**) UV-Vis spectral changes in DosT upon binding O_2_ at various free O_2_ concentrations measured using the optode. **c**) Difference spectra showing spectral changes when DosT binds O_2_. **d**) O_2_ affinity plots for DosT and DosS proteins in dark blue and light blue, respectively (n=3). Gray dashed and dotted lines show the O_2_ concentrations at which NRP-1 and NRP-2 stages are triggered in *Mtb*, respectively. O_2_ concentrations relevant to normoxia, micro-oxia, and nano-oxia are also depicted on the x-axis.

The role of DosT and DosS as two distinct O_2_ sensors in the dormancy pathway of *Mtb* has received considerable attention and has been extensively discussed in literature [3,4,8]. Some studies suggest that the presence of these two sensors aligns with the Wayne’s hypoxia model, which proposes two stages of non-replicating persistence — NRP-1 and NRP-2 [13–16]. Previous research has reported an O_2_ *K*_d_ value of 26 μM for DosT and a stronger O_2_ *K*_d_ of 3 μM for DosS, indicating that DosT responds to moderate drops in O_2_ levels while DosS responds to deeper hypoxia levels [9]. However, it is noteworthy that NRP-1 occurs at O_2_ levels of 2.5 μM (1% O_2_ with respect to air saturation) and NRP-2 occurs at O_2_ levels of 0.15 μM (0.06% O_2_ with respect to air saturation) [17,18], which are significantly different from the reported *K*_d_ values for DosT and DosS, respectively. This discrepancy raises questions about the direct correlation between the O_2_ sensing capability of these heme-based sensors and the observed stages of non-replicating persistence. Some studies have also suggested that DosT and DosS are switched on much *prior to* and *not during* NRP-1 and NRP-2 stages [14]. We note that, while the O_2_ *K*_d_ value of 3 μM for DosS has been widely referred to in literature, an alternative study has reported a much tighter O_2_ *K*_d_ of 0.58 μM [19], which better aligns with the O_2_ levels for NRP-2. The disparity in reported *K*_d_ values highlights the need for reexamining the O_2_ *K*_d_ values for these heme-based sensors. Ultimately, the O_2_ *K*_d_ value for DosT and DosS are key parameters that relate their biochemical activity to their physiological function and their accurate determination is crucial to elucidate their biological roles.

To gain a better understanding of how the O_2_ affinities of DosT and DosS correlate with *Mtb*’s hypoxia adaptation, we conducted detailed equilibrium and kinetic investigations of O_2_ binding to these heme enzymes. We employed a novel method integrating a UV-Vis spectrometer and an O_2_ optode that enables precise O_2_ *K*_d_ measurements (low nanomolar to high micromolar range) of heme proteins [20]. Briefly, the fraction of O_2_-bound heme is monitored via hypsochromic shifts in the heme Soret band, the free dissolved O_2_ is simultaneously measured *in situ* using an O_2_ optode, and a simple Hill’s fit with n=1 for fraction O_2_-bound protein vs free O_2_ concentration yields the *K*_d_ value. This approach is superior to titration-based approaches that employ total added O_2_ as a measurement parameter which are prone to significant errors from O_2_ escaping into the headspace given the 750-fold [21] higher tendency for O_2_ to exist as gas than dissolved in water. Starting from ferrous DosT with a Soret maximum at 430 nm, adding increasing amounts of O_2_ resulted in systematic hypsochromic shifts with increasing O_2_-bound fractions to a Soret maximum of 414 nm corresponding to the ferrous-oxy DosT (**Fig. 1b**). The O_2_-bound DosT fractions are calculated from delta absorbance curves (**Fig. 1c**) as a normalized sum of the magnitude change in absorbances at 410 and 434 nm. A Hill’s fit with n=1 for the O_2_-bound fraction vs the corresponding free O_2_ measured using the optode gives a *K*_d_ value of 3.3 ± 1 μM for DosT (dark blue curve, **Fig. 1d**) which is 8-fold tighter than what has been previously reported [9]. We also note that this value correlates well with O_2_ levels for the onset of NRP-1 (dashed gray line, **Fig. 1d**) as per the Wayne’s model of hypoxia adaptation in *Mtb*. Next, we measured O_2_ affinity of DosS using the same method (**Fig. S3**), and we determined a *K*_d_ value of 0.46 ± 0.08 μM for DosS (light blue curve, **Fig. 1d**). This value is 7-fold tighter than the widely referred O_2_ *K*_d_ value of 3.3 μM for DosS reported by Sousa *et al*. [9], but matches well with the *K*_d_ of 0.58 μM reported by Ioanoviciu *et al* [19]. Again, our measured O_2_ *K*_d_ value of 0.46 μM for DosS correlates well with O_2_ levels that drive the onset of NRP-2 (dotted gray line, **Fig. 1d**). In all, our O_2_ affinity measurements for both DosT and DosS reveal that their respective *K*_d_ values lie well within the nano-oxic regime (O_2_ levels of 0.13 - 6 μM) that are typical for TB granulomas [22]. Next, we conducted stopped-flow kinetic investigations of O_2_ binding to DosT and DosS to determine their ligand association and dissociation rates (**Fig. S4-5**). Upon reacting ferrous DosT with 10 μM O_2_, we observe a rather slow association rate (k_on_ = 0.84 μM^-1^s^-1^) that equilibrates to about 60% ferrous-oxy DosT in the hundred millisecond timescale (dark blue curve, **Fig. 2**). Upon reacting ferrous DosS with 10 μM O_2_, we note a faster association rate (k_on_ = 7.2 μM^-1^s^-1^) that goes to near completion in the tens of millisecond timescale (light blue curve, **Fig. 2**). We note that these k_on_ values match well with previous reports by Sousa *et al*. and Ioanoviciu *et al* [9,19]. Based on k_on_ values from these kinetic investigations and the *K*_d_ values from equilibrium affinity measurements, we calculate the dissociation rate constant (k_off_) for DosT and DosS as 2.8 s^-1^ and 3.3 s^-1^, respectively.

**Figure 2.**
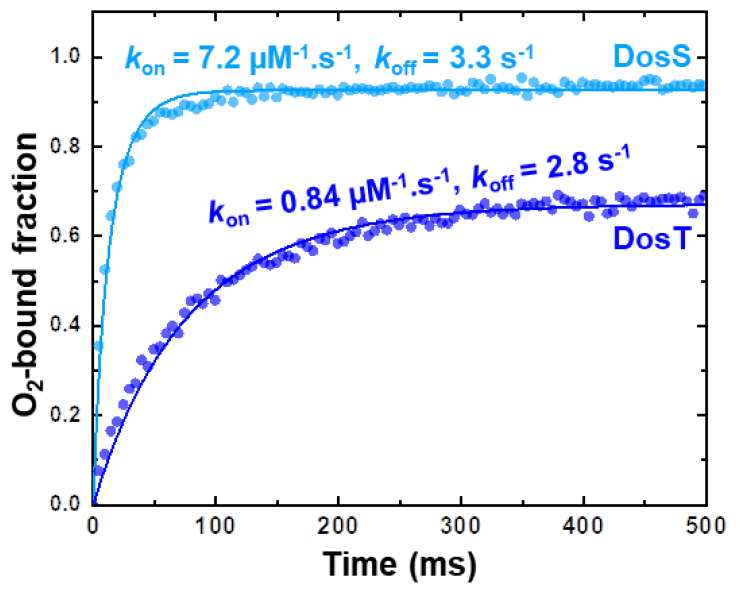
O_2_ binding kinetics for DosT and DosS proteins shown in dark blue and light blue, respectively (n=3).

In summary, our studies rectify a substantial mischaracterization and misconception regarding the O_2_ affinities of DosT and DosS — two important heme-based O_2_ sensors responsible for hypoxia adaptation and dormancy signaling in *Mtb*. We note that the O_2_ *K*_d_ value of 3.3 μM for DosT is 8-fold tighter than previous reports and correlates with O_2_ levels at the onset of NRP-1 (∼2.5 μM O_2_) indicating its role in early adaptation to hypoxic conditions. Given that DosT is constitutively expressed in *Mtb* and not a part of the DosR regulon [4], its solo activation also represents a reversible regime wherein the bacteria can transition back to its active form. Activation of the DosR regulon by DosT during NRP-1 leads to the expression of DosS (part of DosR regulon [4]) with an O_2_ *K*_d_ value of 0.46 μM which transitions to its active form as O_2_ concentrations drop down to NRP-2 levels (∼0.15 μM O_2_). In this regime, activation of DosR regulon by DosS leads to further expression of active DosS, thereby reinforcing dormancy transition and marking the onset of NRP-2 stage in *Mtb*.

## Authorship and Contribution Statement

***Elizabeth A. Apiche***: Investigation – Expression, purification, and characterization of DosT, Writing – original draft; ***Eaindra Yee***: Investigation – Expression, purification, and characterization of DosS, Writing – original draft; ***Anoop Rama Damodaran***: Equilibrium and kinetic investigations of DosS and DosT, Writing – original draft, Supervision, Conceptualization; ***Ambika Bhagi-Damodaran***: Equilibrium and kinetic investigations of DosS and DosT, Writing – original draft, Supervision, Funding acquisition

## Data availability

Data will be made available on request.

## Acknowledgements

This work was supported by the Regents of the University of Minnesota and NIH NIGMS grant # R35GM138277. E.A.A. and E.Y. acknowledge the support of Lester C. and Joan M. Krogh Endowed Excellence Fellowship and PPG Inc. Endowed Graduate Student Excellence Fellowship from the Department of Chemistry, University of Minnesota. The authors thank Prof. Robert Abramovitch (Michigan State University) for the DosT plasmid.

