## Supplementary Information for "Oxygen affinities of DosT and DosS sensor kinases with implications for hypoxia adaptation in *Mycobacterium tuberculosis*"

Elizabeth A. Apiche[¶§], Eaindra Yee[¶§], Anoop Rama Damodaran[¶*], Ambika Bhagi-Damodaran[¶*]
[¶]Department of Chemistry, University of Minnesota, Minneapolis, MN 5545

[§]These authors contributed equally

**Materials and Methods:**

Expression and purification of WT full-length DosT:

Wild-type full-length DosT was expressed largely following the protocol reported previously [1]. Briefly, transformation of pET15b vector with WT FL DosT was performed in conjunction with pTGroE in *E. coli* BL21-Gold (DE3) competent cells. Primary cultures were inoculated overnight at 37°C in four 25-mL LB medium with appropriate antibiotics (50 µg/mL kanamycin and 37 µg/mL chloramphenicol). ZYP (10 g tryptone, 5 g yeast, 923 mL water)-5052 (0.5 % glycerol, 0.05% glucose, 0.2% a-lactose) auto-induction medium (1 L) containing 37 µg/mL chloramphenicol and 100 µg/mL kanamycin was used for secondary cultures. Cultures were grown at 37°C for 4 hours and 30 mg/L hemin was added to each flask. Temperature was lowered to 18°C and cultures were grown for additional 24 hours. Cells were harvested by centrifuging at 8,000 rpm at 4°C, flash frozen, and stored in the -20°C freezer.

Full-length DosT was purified following DosS purification protocol with a few notable differences. Lysis buffer used to lyse the cells was 50 mM $NaH_2PO_4$, 200 mM NaCl, 10% glycerol, 1% Triton X-100, 1 mM PMSF, pH = 7.5. After IMAC, eluted DosT fractions were collected and dialyzed in 20 mM Tris-HCl buffer (pH 7.5) overnight using 25.5 mm spectra/Por molecular porous membrane tubing. Recovered protein was buffer-exchanged into storage buffer (50 mM Tris-HCl, 100 mM NaCl, 5% glycerol pH 7.5), concentrated, and stored in -80°C until usage. SDS-PAGE analysis was performed to assess protein purity (**Fig. S1**).

Expression and purification of WT full-length DosS:

The full-length DosS contained in the vector pET23a(+) was co-transformed with GroES/EL plasmid into BL21-Gold (DE3) competent cells (Agilent) per commercial protocol. The cells were then plated onto an LB-agar plate supplemented with ampicillin (100 µg/mL) and chloramphenicol (37 µg/mL) and grown overnight at 37°C. The overnight colonies were then inoculated into TB medium supplemented with the same concentrations of the same antibiotics and grown overnight at 37°C with shaking (200 rpm). 10 mL of the overnight culture was used to inoculate 1 L TB medium containing 100 µg/mL ampicillin and 37 µg/mL chloramphenicol. The cells were grown to optical density, $OD_{600}$, of 0.6-0.8 before adding hemin and zinc (II) acetate to the final concentrations of 0.5 mM and 40 µM, respectively. The cells were finally induced with 0.75 mM IPTG at 16°C for 48 hours. After overexpression, the cells were harvested by centrifugation at 8,000 rpm at 4°C, flash-frozen, and stored at -20°C until usage.

To purify full-length DosS, the collected cell pellet was resuspended in lysis buffer (50 mM $NaH_2PO_4$, 250 mM NaCl, 10% glycerol, 1% Triton X-100, 0.1 mg/mL DNase, 0.1 mg/mL RNase, 1 mg/mL lysozyme, 1 tablet/50 mL Pierce™ Protease Inhibitor, pH = 7.5), sonicated, centrifuged (20,000 rpm, 4°C), and filtered (using MCE Membrane Filter, 0.22 µm) before loaded onto 5-mL Histrap FF column on an AKTA Start protein purification system. After sample application, the unbound proteins were washed out with 10 CV of IMAC buffer A (50 mM $NaH_2PO_4$, 500 mM NaCl, 20 mM imidazole, 10 % glycerol, pH 7.5) and the protein of interest was eluted using a two-step gradient method: 10% IMAC buffer B (50 mM $NaH_2PO_4$, 500 mM NaCl, 400 mM imidazole, 10 % glycerol, pH 7.5) for 7 CV, then 100% buffer B for 5 CV. Both the wash out unbound and the elution steps were performed at the flow rate of 2 mL/min. The fractions from the last elution step were pooled and buffer-exchanged with ~90 mL DEAE buffer A (50 mM Tris-HCl, 1 mM EDTA, 5% glycerol, pH = 8.00) using an Amicon Ultra-15 centrifugal unit (50 kDa MWCO). The buffer-exchanged solution was subsequently applied onto a 5-mL HiTrap DEAE FF on AKTA Start system. After sample application, the column was washed with 7 CV of DEAE buffer A and the full-length DosS was eluted using a linear-gradient method: 0-100% of DEAE buffer B (50 mM Tris-HCl, 500 mM NaCl, 1 mM EDTA, 5% glycerol, pH = 8.00) for 12 CV. Fractions that corresponded to the peak of UV chromatogram were collected, concentrated, and the purity of the concentrated proteins were determined by SDS-PAGE (**Fig. S1**) before being aliquoted, flash-frozen, and stored in -80°C until usage.

Hemochromogen assay:

To determine the extinction coefficients of full-length DosT and DosS, a pyridine hemochromogen assay was performed according to the published protocol [2]. Briefly, 100 µL of ferric protein was mixed with 80 µL 5M NaOH and 800 µL pyridine. The fully oxidized spectrum was recorded before adding 10 uL of 0.5 M sodium dithionite solution to obtain the reduced mixture. The absorbance value at 557 nm was recorded and the molar absorptivity of pyridine-heme *b* hemochromogen (34.7$mM^{-1}cm^{-1}$) [3] was used to finally determine the extinction coefficients of full-length ferric DosS and ferrous-$O_2$ DosT to be 145 $mM^{-1}cm^{-1}$ and 172 $mM^{-1}cm^{-1}$, respectively (**Fig. S2**).

Measurement of $O_2$ affinity values for DosT and DosS proteins:

The $O_2$ affinity measurement and data analysis for DosT and DosS proteins were performed as described step-by-step in a methods paper published by our lab previously [4].

Measurement of $O_2$ association and dissociation constants for DosT and DosS proteins:

Experiments were performed inside an anaerobic glove bag using a stopped-flow UV-Vis observation cell connected to broad band light source and a fast photodiode array detector. Ferrous DosT and DosS were prepared by reduction with 10 equivalents dithionite followed by removal of dithionite by flowing through a PD-10 column and concentration via MWCO = 10 kDa centricons. Two-syringe mixing was employed to mix equal volumes of 7.4 µM of ferrous DosT or 9 µM ferrous DosS with buffer containing 20 µM $O_2$ (measured using an $O_2$ optode). All reported data sets originally consisted of 500 spectra collected over 500 ms using linear sampling. A water bath connected to the syringe compartment and the observation cell of the stopped-flow instrument was set to 22ºC and provided temperature control. The kinetic traces for $O_2$-bound protein (y) were obtained by fitting the data to the following equation describing second-order kinetics: $\frac{d[y]}{dt} = k_{on}.([P]-y).([O_2]-y)$ - $K_D.k_{on}.y$ where [P] is the total protein concentration, $K_D$ is the equilibrium dissociation constant measured via equilibrium $O_2$ titration studies, and $k_{on}$ is the association rate constant. The fitting procedure yielded $k_{on}$, and the dissociation rate constant ($k_{off}$) was calculated using the relationship: $k_{off} = K_D.k_{on}$

**SI Figures**

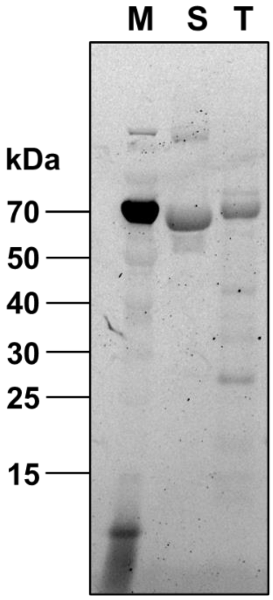


**Figure S1:** SDS-PAGE gel of full-length *Mtb* DosS (lane indicated as S) and *Mtb* DosT (lane indicated as T) investigated in this study. Expected MW of DosT and DosS are 64.44 kDa and 64.05 kDa, respectively. The MW marker lane is indicated as M.

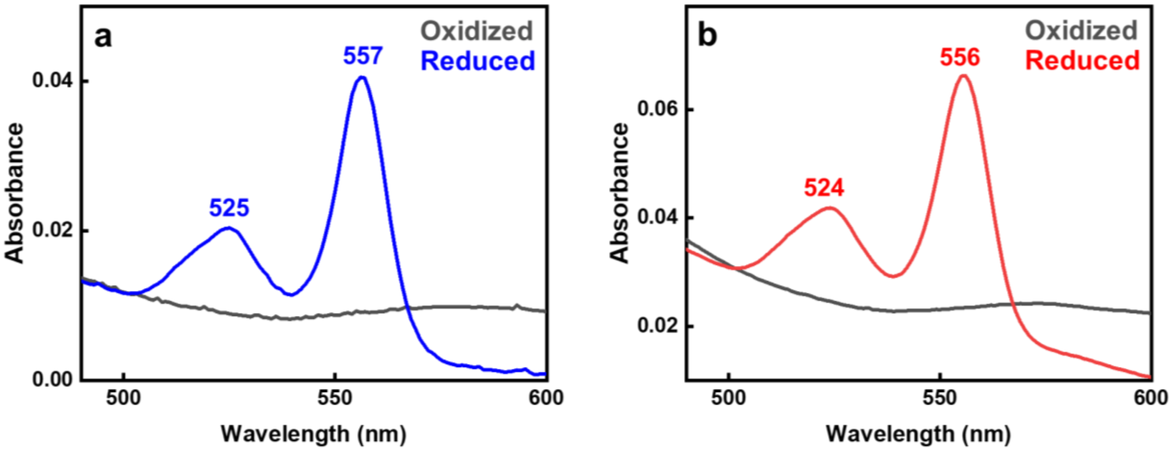


**Figure S2:** Pyridine hemochromogen assays were performed on full-length *Mtb* (a) DosS and (b) DosT to find the extinction coefficients of the proteins based on the spectral features. Spectra of reduced (blue for DosS and red for DosT) and oxidized (dark grey) pyridine hemochromogen with notable peaks shown. The Soret extinction coefficients were calculated as 145 ± 2 $mM^{-1}cm^{-1}$ (n = 3) and 172 ±19 $mM^{-1}cm^{-1}$ (n = 3) for ferric DosS and ferrous-$O_2$ DosT, respectively.

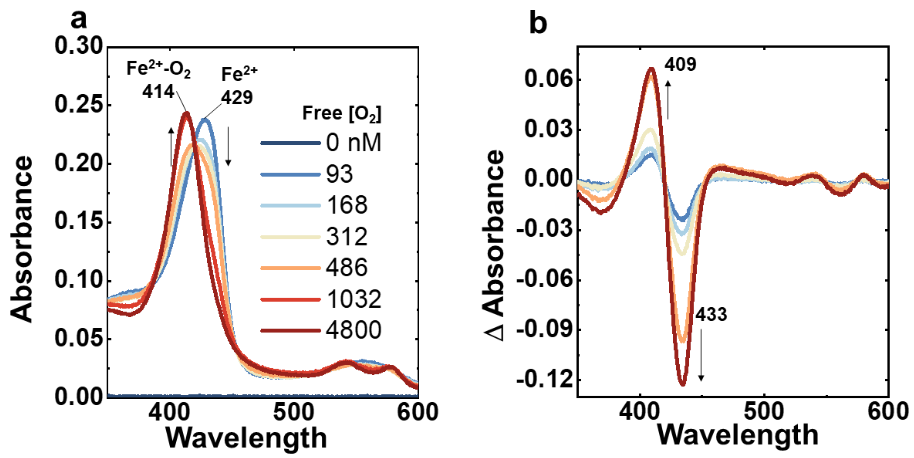


**Figure S3:** a) UV-Vis spectral changes in DosS upon binding $O_2$ at various free $O_2$ concentrations measured using the optode. b) Difference spectra showing spectral changes in DosS when it binds $O_2$.

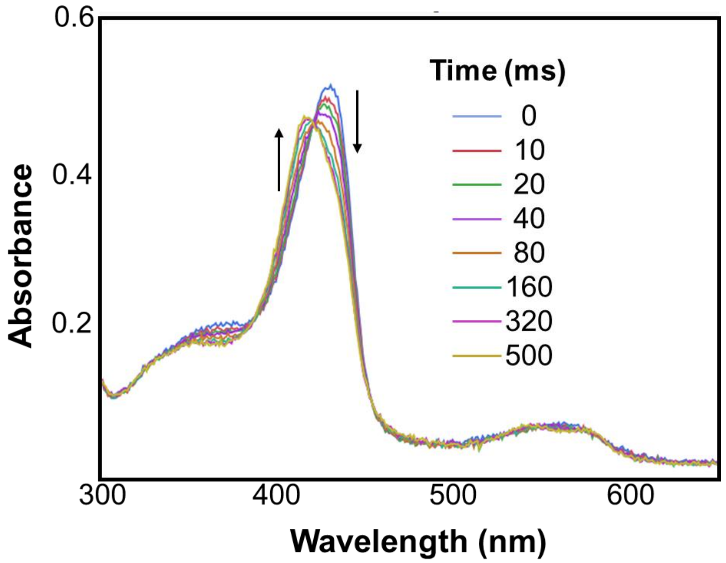


**Figure S4**: UV-Vis spectral changes in DosT upon binding $O_2$ at specific time points as observed in stopped-flow kinetic studies.

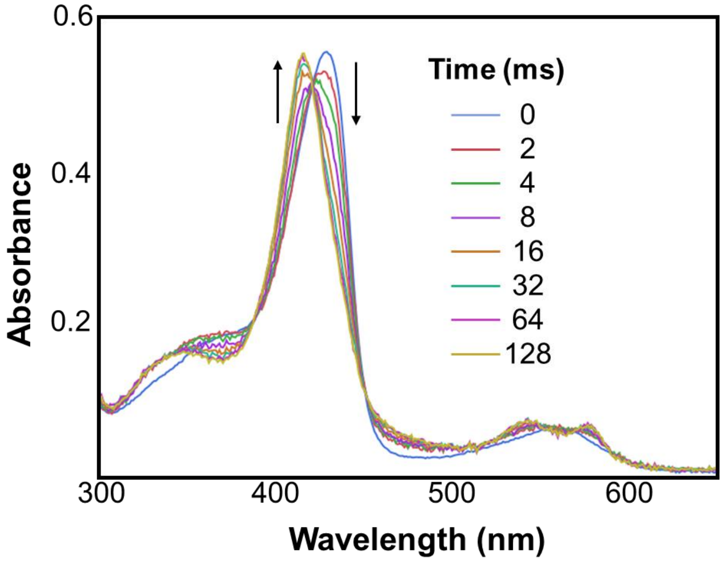


**Figure S5**: UV-Vis spectral changes in DosS upon binding $O_2$ at specific time points as observed in stopped-flow kinetic studies.
